# COVID-19 CG: Tracking SARS-CoV-2 mutations by locations and dates of interest

**DOI:** 10.1101/2020.09.23.310565

**Authors:** Albert Tian Chen, Kevin Altschuler, Shing Hei Zhan, Yujia Alina Chan, Benjamin E. Deverman

**Affiliations:** Stanley Center for Psychiatric Research, Broad Institute of MIT and Harvard, Cambridge, MA, United States of America; Department of Zoology & Biodiversity Research Centre, the University of British Columbia, Vancouver BC, Canada

## Abstract

COVID-19 CG is an open resource for tracking SARS-CoV-2 single-nucleotide variations (SNVs) and lineages while filtering by location, date, gene, and mutation of interest. COVID-19 CG provides significant time, labor, and cost-saving utility to diverse projects on SARS-CoV-2 transmission, evolution, emergence, immune interactions, diagnostics, therapeutics, vaccines, and intervention tracking. Here, we describe case studies in which users can interrogate (1) SNVs in the SARS-CoV-2 Spike receptor binding domain (RBD) across different geographic regions to inform the design and testing of therapeutics, (2) SNVs that may impact the sensitivity of commonly used diagnostic primers, and (3) the recent emergence of a dominant lineage harboring an S477N RBD mutation in Australia. To accelerate COVID-19 research and public health efforts, COVID-19 CG will be continually upgraded with new features for users to quickly and reliably pinpoint mutations as the virus evolves throughout the pandemic and in response to therapeutic and public health interventions.

## Introduction

Since the beginning of the pandemic, SARS-CoV-2 genomic data has been accumulating at an unprecedented rate (90,000+ virus genomes as of September, 2020 on the GISAID database) (Elbe and Buckland-Merrett, 2017; Shu and McCauley, 2017). Numerous countries have mobilized to sequence thousands of SARS-CoV-2 genomes upon the occurrence of local outbreaks, collectively and consistently contributing more than 10,000 genomes per month (**Figure S1A, B**). It is important to note that, despite the slow accumulation of potentially functional (nonsynonymous) mutations, there has been a steady increase in the number of variants with more than 6 nonsynonymous mutations compared to the WIV04 reference, an early isolate of SARS-CoV-2 that was collected in Wuhan in December, 2019 (**Figure S1C**). To evaluate the outcomes of anti-COVID-19 measures and detect keystone events of virus evolution, it is important to track changes in SARS-CoV-2 mutation and population dynamics in a location and date-specific manner. Indeed, several countries and the National Institutes of Health (NIH) have recognized how critical it is to collect SARS-CoV-2 genomic data to support contact tracing efforts and to inform public health decisions – these are paramount to the re-opening of countries and inter-regional travel (Collins 2020; Rockett et al. 2020; Oude Munnink, et al. 2020; Gudbjartsson et al. 2020; Pybus et al. 2020). Yet, the quantity and complexity of SARS-CoV-2 genomic data (and metadata) make it challenging and costly for the majority of scientists to stay abreast of SARS-CoV-2 mutations in a way that is meaningful to their specific research goals. Currently, each group or organization has to independently expend labor, computing costs, and, most importantly, time to curate and analyze the genomic data from GISAID before they can generate specific hypotheses about SARS-CoV-2 lineages and mutations in their population(s) of interest.

## Results

To address this challenge, we built COVID-19 CoV Genetics (COVID-19 CG, covidcg.org), a performant, interactive, and fully-scalable web application that tracks SARS-CoV-2 single-nucleotide variants (SNVs) and lineages without sub-sampling. COVID-19 CG is a free, open access interface that allows users to adapt analyses according to their dates and locations of interest (**Figure 1A**,**B**; data processing workflow in **Figure S2**). Users can also select and compare trends in SARS-CoV-2 lineage or SNV frequency across multiple locations (**Figure 1C**) as we will demonstrate using case studies. COVID-19 CG provides functionalities that, to the best of our knowledge, cannot be found in other existing public browsers, and was designed to empower these specific user groups:

**Figure 1.**
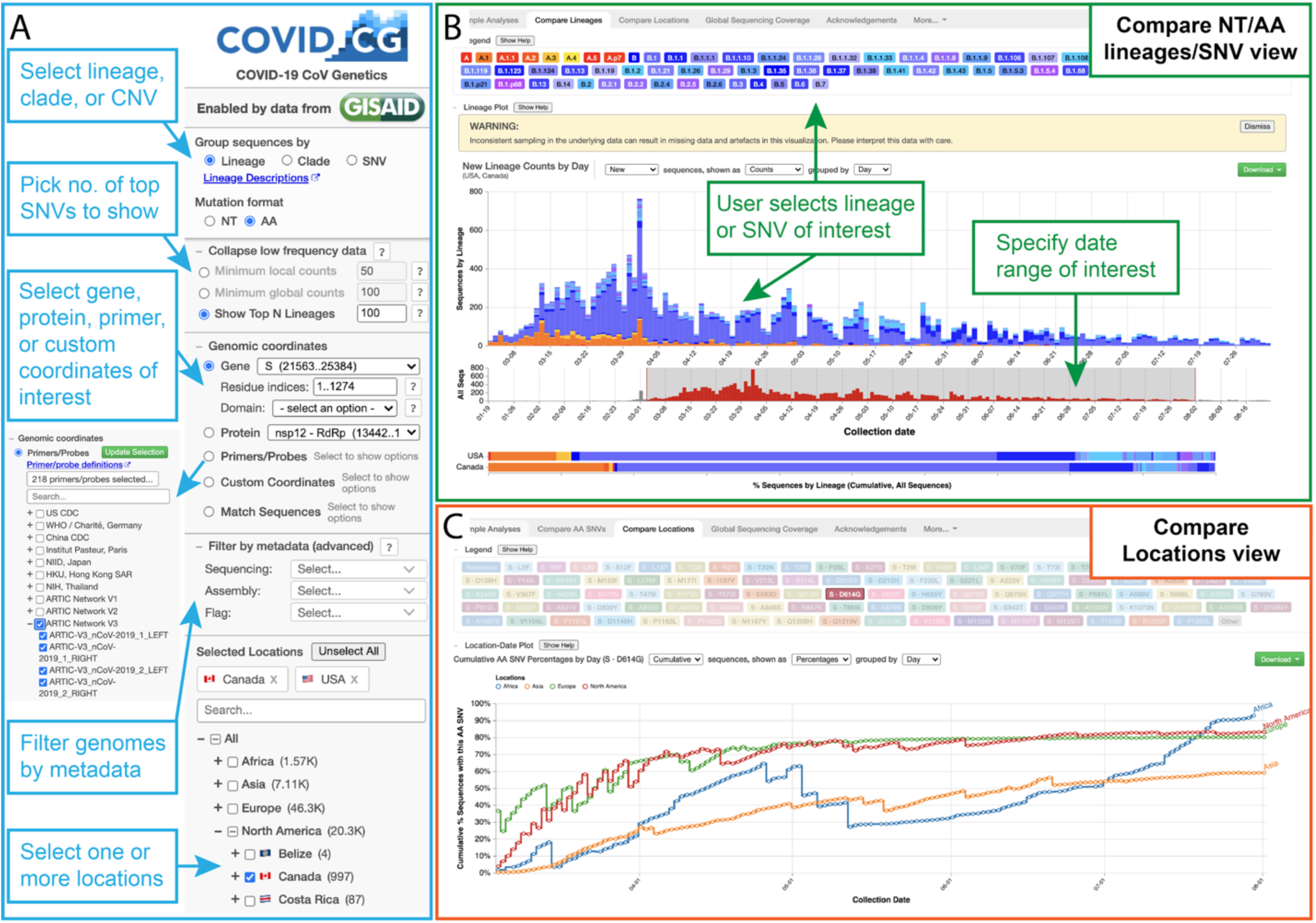
The COVID-19 CG (https://covidcg.org) browser interface. **(A)** Users can select SARS-CoV-2 genomes according to lineage, clade, or SNV, virus gene or protein, and location(s). Genomes can also be filtered by metadata, and specifically analyzed at genomic coordinates of interest, such as the target sites of 665 commonly used diagnostic primers and probes. **(B)** In the “Compare lineages or SNVs” tab, users can visualize SARS-CoV-2 lineages or SNVs by location, define their date range of interest, and see the corresponding SNVs at the nucleotide or amino acid level. **(C)** In the “Compare locations” tab, users can compare the frequencies of specific lineages or SNVs in multiple locations over time.

***Vaccine and therapeutics developers*** can inform the design and testing of their vaccine, antibody, or small molecule by using COVID-19 CG to rapidly identify all of the variants in their targeted SARS-CoV-2 protein or antigen, alongside the frequency of each variant in their geographic location(s) of interest. Scientists can use COVID-19 CG to generate hypotheses and experimentally determine whether the variants present in the location of vaccine/therapeutic implementation may impact their product-specific interaction interface or antigen.

### Case study of SNVs in the receptor binding domain (RBD) of the SARS-CoV-2 Spike

Analyzing SNVs by geography and time is critical as the frequency of each SNV may vary significantly across different regions over time. For instance, an S477N mutation in the RBD has become dominant in Oceania (57.5% of Oceanian genotypes, all time) although it constitutes only 4.36% of SARS-CoV-2 genotypes globally and has not been detected in Africa, Asia, or South America (**Figure 2A**). SNV frequency in a given region can also shift over time, e.g., an RBD N439K mutation not found in Ireland prior to July is now present in 79.5% of the genomes collected mid-July through August (**Figure 2B**). Another rare RBD S477N mutation, which was found in only 1.05% of the Australian SARS-CoV-2 sequences before June, now constitutes more than 90% of the sequenced June through August isolates (**Figure 2C**). This geographical and temporal variation is important to incorporate into the design and testing of therapeutic antibodies (such as those under development as therapeutics by Regeneron that specifically target the SARS-CoV-2 Spike RBD), as well as mRNA or recombinant protein-based vaccines. This will help to assure developers of the efficacy of their therapeutics and vaccines against the SARS-CoV-2 variants that are present in the intended location of implementation.

**Figure 2.**
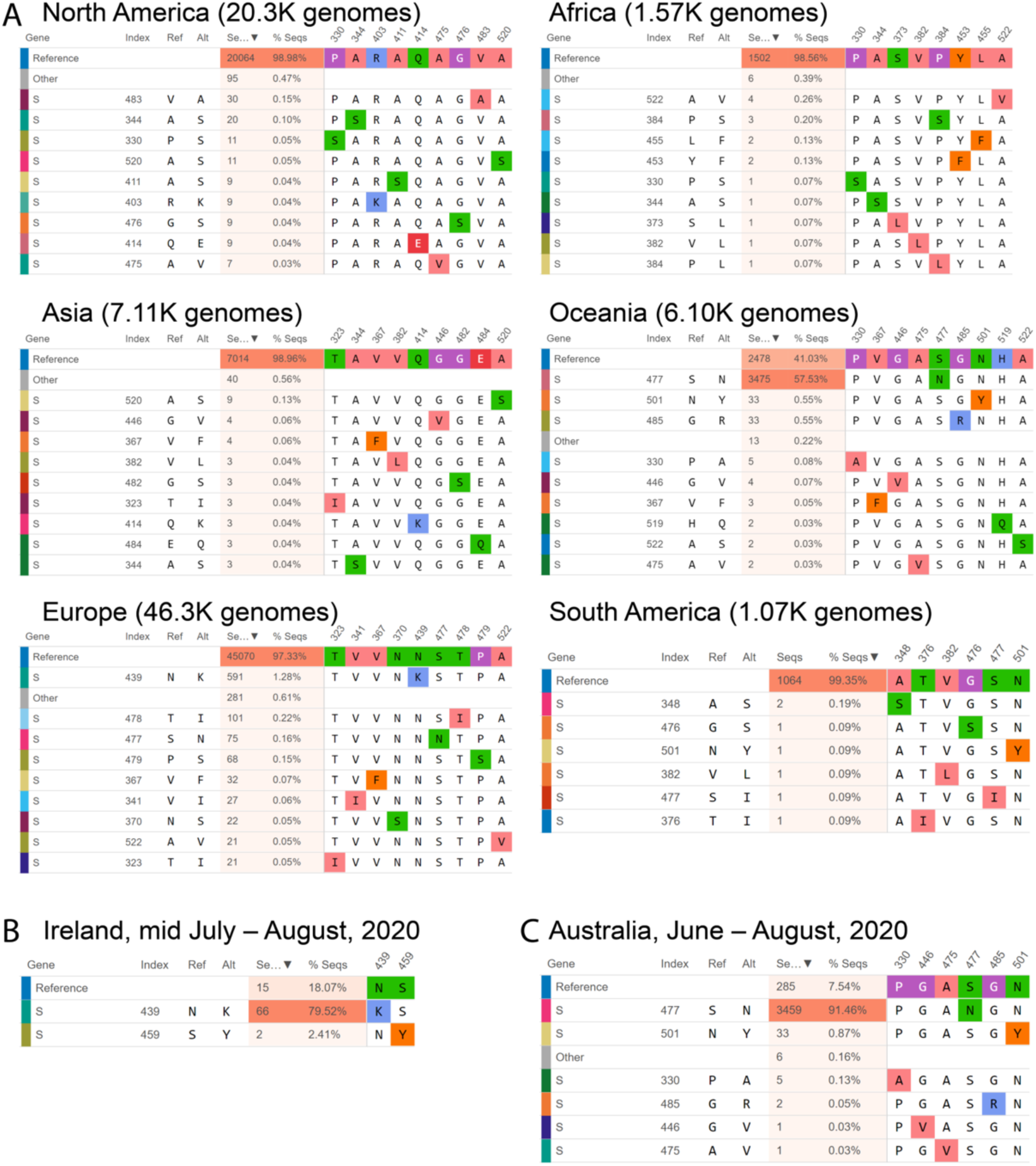
Mutational frequencies in the SARS-CoV-2 Spike receptor binding domain (RBD) across geographical location and time. Screen captures from the Compare AA SNVs tab are shown. **(A)** The top 10 RBD SNVs alongside the number of high quality sequences available on GISAID are shown for each region. **(B)** The top RBD SNVs for Ireland between mid July and August, 2020 are shown. The S439N mutant had not been previously detected in Ireland. **(C)** The top RBD SNVs for Australia between June and August, 2020 are shown. The S477N mutant constituted only 1.05% of the Australian SARS-CoV-2 genomes on GISAID prior to June.

In addition, COVID-19 CG can be harnessed to track changes in SARS-CoV-2 evolution post-implementation of therapeutics and vaccines. It will be crucial to watch for rare escape variants that could resist drug- or immune-based interventions to eventually become the dominant SARS-CoV-2 variant in the community. This need was particularly emphasized by a Regeneron study that demonstrated that single amino acid variants could evolve rapidly in the SARS-CoV-2 Spike to ablate binding to antibodies that had been previously selected for their ability to neutralize all known RBD variants; these amino acid variations were found either inside or outside of the targeted RBD region, and some are already present at low frequency among human isolates globally (Baum et al., 2020). The authors, Baum et al., suggested that these rare escape variants could be selected under the pressure of single antibody treatment, and, therefore, advocated for the application of cocktails of antibodies that bind to different epitopes to minimize SARS-CoV-2 mutational escape. A recent study by Greaney et al. generated high-resolution ‘escape maps’ delineating RBD mutations that could potentially result in virus escape from neutralization by ten different human antibodies (Greaney et al., 2020). Based on lessons learnt from the rise of multidrug resistant bacteria and cancer cells, it will be of the utmost importance to continue tracking SARS-CoV-2 evolution even when multiple vaccines and therapeutics are implemented in a given human population.

***Diagnostics developers*** can evaluate their probe, primer, or point-of-care diagnostic according to user-defined regional and temporal SARS-CoV-2 genomic variation. More than 665 established primers/probes are built into COVID-19 CG, and new diagnostics will be continually incorporated into the browser. Users can also input custom coordinates or sequences to evaluate their own target sequences and design new diagnostics.

### Case study of SNVs that could impact the sensitivity of diagnostic primers

A recent preprint alerted us to the finding that a common G29140T SNV, found in 22.3% of the study’s samples from Madera County, California, was adversely affecting SARS-CoV-2 detection by the NIID_2019-nCoV_N_F2 diagnostic primer used at their sequencing center; the single SNV caused a ∼30-fold drop in the quantity of amplicon produced by the NIID_2019-nCov_N_F2/R2 primer pair (Vanaerschot et al., 2020). We used COVID-19 CG to detect other SNVs that could impact the use of this primer pair, discovering that there are SARS-CoV-2 variants in several countries with a different C29144T mutation at the very 3’ end of the same NIID_2019-nCoV_N_F2 primer (**Figure 3A**). As the authors of the preprint, Vanaerschot et al., noted, SNVs could impact assay accuracy if diagnostic primers and probes are also being used to quantify viral loads in patients. We found that at least ten other primer pairs could potentially be at risk in different geographical regions due to SNVs that appear proximal to the 3’ ends of primers (**Figure 3B–K**): China-CDC-N-F and R; NIH, Thailand, WH-NIC N-F; US CDC 2019-nCoV-N1-R; US CDC 2019-nCoV-N2-F; ARTIC-V3_nCoV-2019_11_RIGHT; ARTIC-V3_nCoV-2019_13_LEFT; ARTIC-V3_nCoV-2019_34_LEFT; ARTIC-V3_nCoV-2019_39_LEFT (note that the ARTIC primers are used for nanopore sequencing) (Tyson et al., 2020); WHO N_Sarbarco_R1; and Institut Pasteur, Paris 12759Rv. We advocate that labs and clinics use COVID-19 CG (https://covidcg.org) to check their most commonly used primers and probes against the SARS-CoV-2 sequences that are prevalent in their geographic regions.

**Figure 3.**
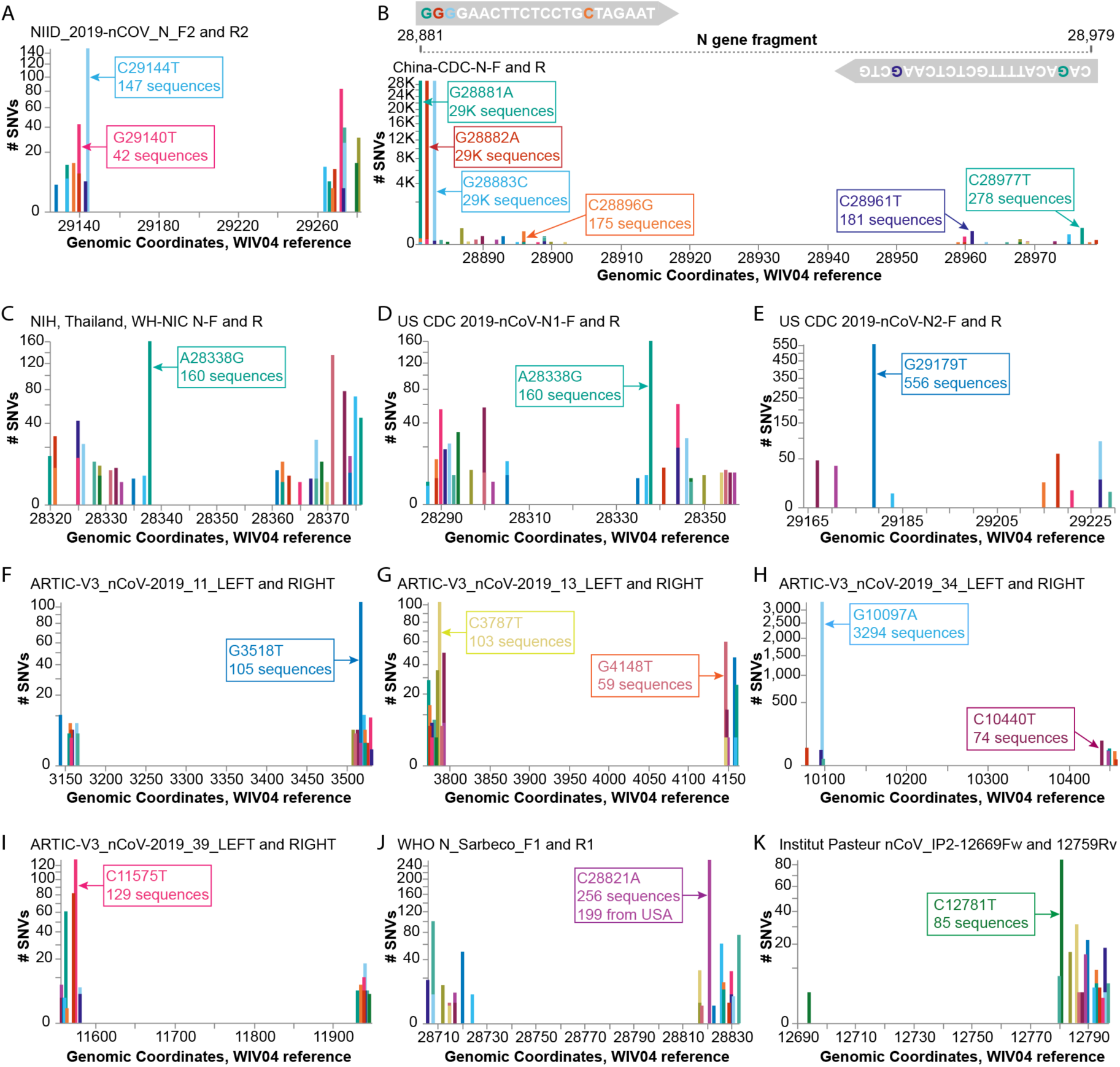
Investigating diagnostic-targeted regions of the SARS-CoV-2 genome for SNVs that could impact primer/probe sensitivity. Images were downloaded from the Compare NT SNVs tab. Labels for specific mutations were added. Primer pairs that contain at least one primer with potentially impactful SNVs near the 3’ end are shown. None of the 11 primer pairs shown here were designed with degenerate bases. (**A)** The G29140T has been demonstrated to impact the NIID_2019-nCOV_N_F2 primer sensitivity. **(B-K)** Primer pairs affected by SNVs with a global frequency of more than 80 instances are shown. **(B)** As an example, majors SNVs are colored accordingly in the China-CDC-N-F and R (forward and reverse) primers.

***Researchers and public health professionals*** can use COVID-19 CG to gain insights as to how the virus is evolving in a given population over time (e.g., in which genes are mutations occurring, and do these lead to structural or phenotypic changes?). For example, users can track D614G distributions across any region of interest over time. **Figure 4** shows a variety of different D614G population dynamics in different areas. Nonetheless, we strongly caution against inferring (i) chains or directionality of transmission and (ii) changes in the transmissibility of any SARS-CoV-2 SNV based on population dynamics alone. Inconsistent sampling, sampling biases, differences in founder host population traits (even median patient age), superspreading events, regionally and temporally differential travel restrictions, and numerous other factors instead of virus biological differences can influence the global distribution of SNVs (Grubaugh et al., 2020).

**Figure 4.**
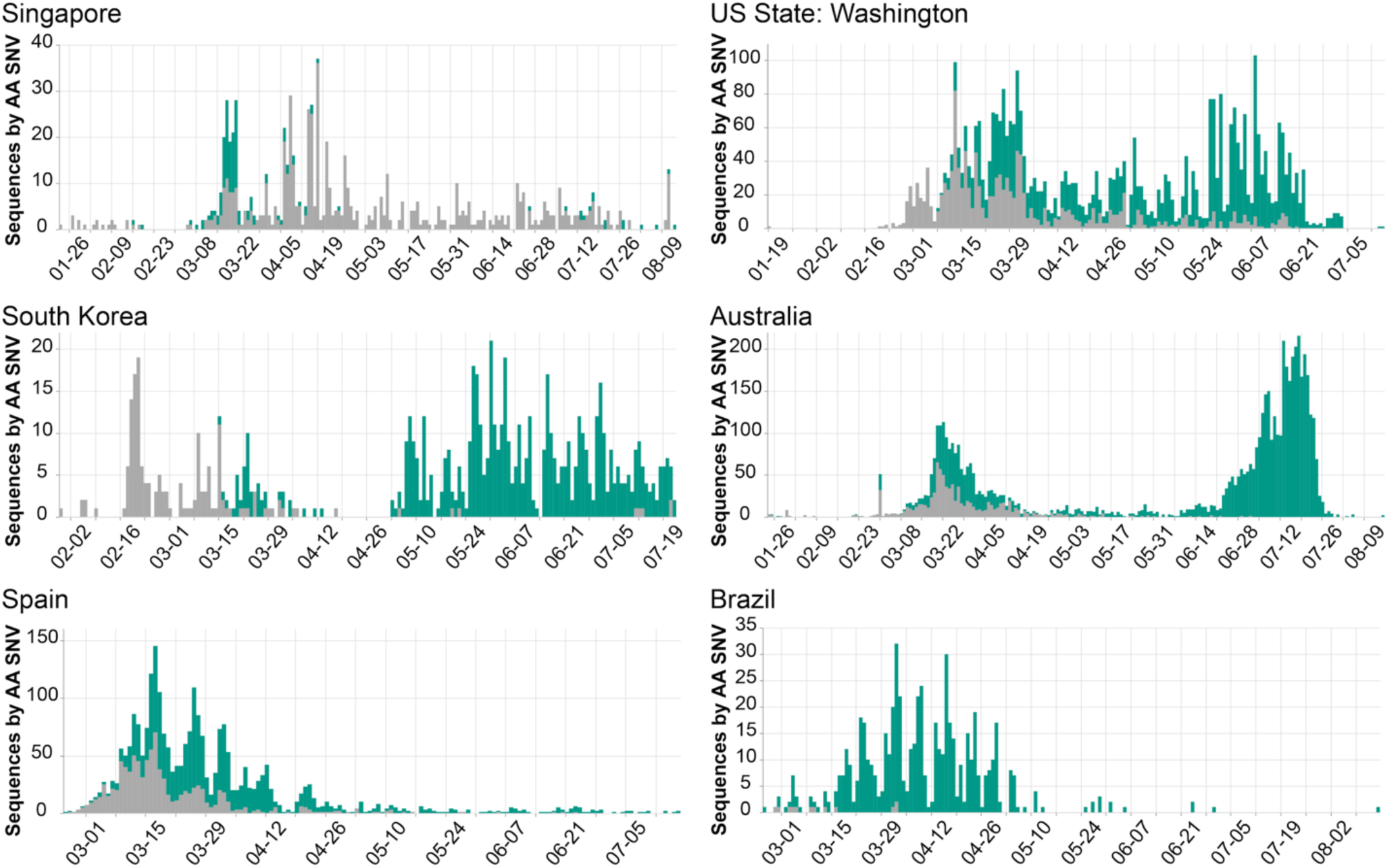
Population dynamics of Spike D614G in different regions. Images were downloaded from the Compare Lineages tab of covidcg.org: The Spike D614 variants are shown in grey, and the G614 variants are shown in green. Plots displaying different population dynamics were deliberately selected. Time is shown on the horizontal axis and the number of sequences is shown on the vertical axis; these differ per country depending on when and how many samples were collected and whether the sequences were deposited onto GISAID by August 31, 2020.

### Case study of Australia’s new dominant SARS-CoV-2 variant

We discovered that the SARS-CoV-2 Spike S477N mutation has become more prevalent in Australia (**Figure 5A**). Globally, the S477N mutation was first detected in a single sample of lineage B.1.1.25 that was collected on January 26, 2020 in Victoria, Australia; this is now the dominant SARS-CoV-2 variant in the region (**Figure 5B, C**). In particular, the set of SNVs that co-occur with the S477N mutation in Australia (all time, as well as prior to May, 2020 before the most recent outbreak) are different from the set of co-occurring SNVs in the United Kingdom (**Figure 5C**) — suggesting that the S477N mutation occurred separately in the Australian and the UK lineages. However, COVID-19 CG only reflects data contributed to GISAID. Variants of interest could be present in other countries, but not yet known to the public because the sequencing centers in those countries have not collected or deposited their data in GISAID. Furthermore, in instances where only a singular, sporadic variant is detected (no sustained transmission), there is also the possibility of sequencing error resulting in incorrect lineage assignment. Due to these caveats, the genetic data must be used in combination with other types of data, such as from contact tracing efforts, before it is possible to draw conclusions about the international transmission of SARS-CoV-2 variants. In the case of the S477N variant that is now dominating in Australia, the sequencing data alone indicate that the local transmission of this variant since January, 2020 in Australia cannot be ruled out.

**Figure 5.**
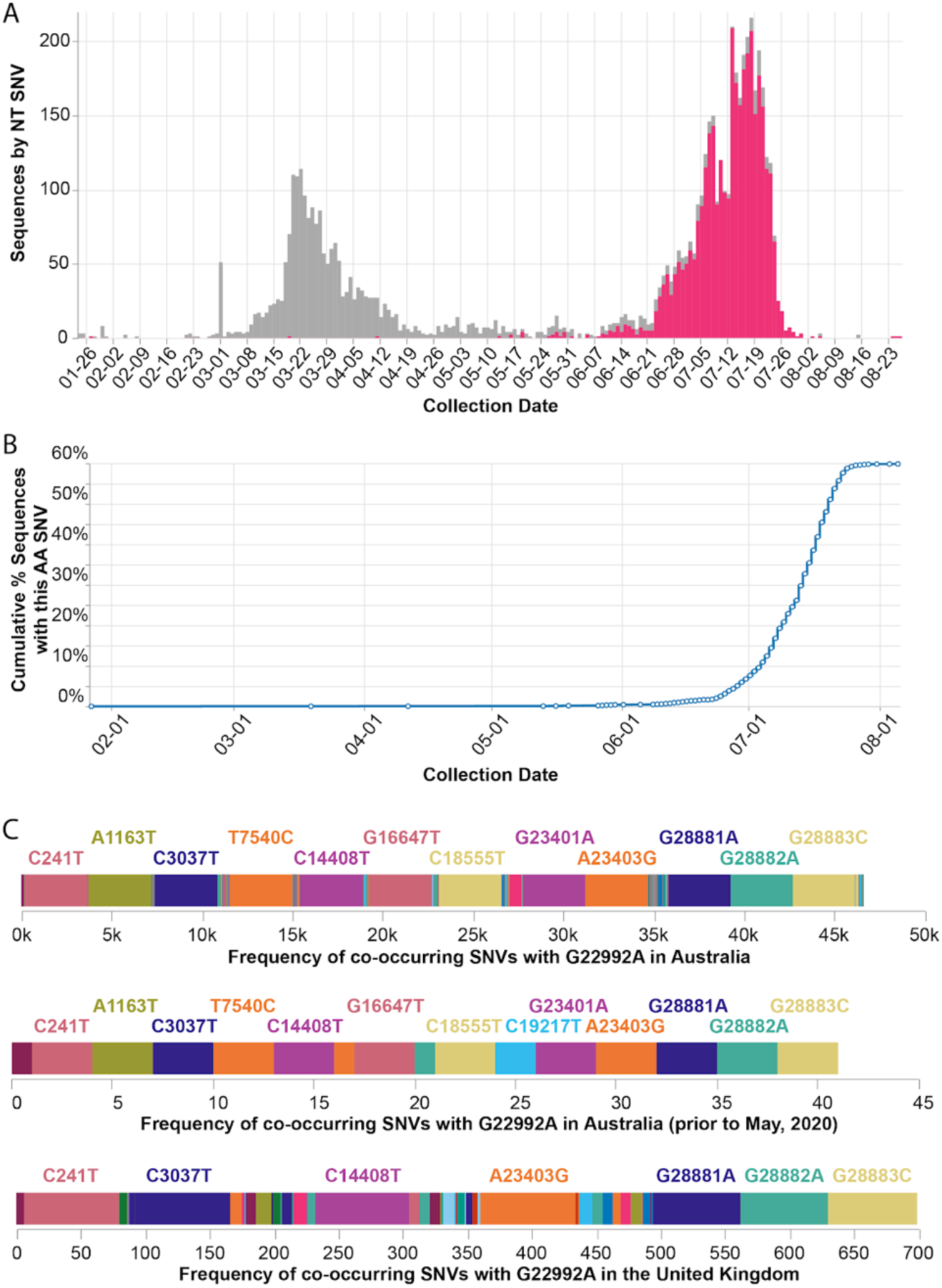
Frequency of the Spike S477N mutation in Australia over time. **(A)** Image downloaded from the Compare NT SNVs tab of covidcg.org: SARS-CoV-2 variants bearing the Spike S477N mutation (also known as the G22992A SNV; depicted in pink), the majority of which lie in the B.1.1.25 lineage, have become the most prevalent form of SARS-CoV-2 in Australia. **(B)** Image downloaded from the Compare Locations tab of covidcg.org: the cumulative percent of sequences carrying the S477N mutation in Australia. **(C)** Images downloaded from the Compare NT SNVs tab of covidcg.org: Co-occurring SNVs of G22992A (Spike S477N) in Australia, all time versus prior to May, 2020, versus in the United Kingdom.

## Discussion

COVID-19 CG (https://covidcg.org) was designed to be modular in order to continually integrate datasets from other COVID-19 initiatives. We anticipate building modules for users to **(1)** map emerging mutations onto structural interaction interfaces of interest (e.g., between virus protein and therapeutic antibodies or host proteins) using existing and future structures on the Protein Data Bank (PDB), **(2)** visualize mutations in isolates of interest in the context of different virus protein phenotypes or mutational constraints of antibody epitopes according to emerging genotype-to-phenotype maps (Greaney et al., 2020; Starr et al., 2020), **(3)** compare SARS-CoV-2 mutations in different host species (e.g., humans versus minks) (Oude Munnink et al., 2020b), **(4)** rapidly determine when and where each lineage or SNV has been detected around the world, and **(5)** overlay important policy events or travel restrictions over time on the lineage or SNV tracker to help guide user date range selection. In addition, as more detailed metadata is generated by COVID-19 studies and initiatives, we will update the application to enable filtering according to patient traits such as gender, age, ethnicity, and medical condition (e.g., symptoms, hospitalization).

COVID-19 CG (https://covidcg.org) was built to help scientists and professionals worldwide, with varying levels of bioinformatics expertise, in their real-time analysis of SARS-CoV-2 genetic data. We hope that COVID-19 CG will also motivate decision makers to sustain or accelerate their sequencing of virus isolates in their geographical area for the purposes of informing vaccine, therapeutics, and policy development. Collecting virus genomic data is particularly relevant to regions that are experiencing increases in COVID-19 cases. If only sparse genomic data are sampled, we risk the late detection of SARS-CoV-2 variants that exhibit enhanced virulence or resistance against therapeutics or vaccination programs in these pandemic hotspots. Furthermore, the widespread implementation of vaccines and antibody therapies could stimulate the emergence and selection of new escape variants (Baum et al., 2020). To compound these risks, SARS-CoV-2 transmission from humans to minks (and back into humans) has already been detected at farms across the Netherlands, Denmark, Spain, and the United States (Oude Munnink et al., 2020b). This process of species crossing, if left unchecked, can result in the emergence of diverse SARS-CoV-2 variants.

Coordinated sequencing and contact tracing efforts (e.g., in the UK, Singapore, the Netherlands, Italy, California, and Australia) emphasize the urgency of establishing open access platforms to evaluate trends in virus introduction into each country or region in real time. Local policymakers, public health researchers, and scientists can use **global** SARS-CoV-2 genetic data, in complementation with contact tracing data, to better understand which lineages were imported into their region (from which potential international locations), whether these were introduced multiple times, and if particular lineages are dying out or persisting. Labs in numerous countries are already making incredible efforts to sequence the SARS-CoV-2 variants circulating in their local populations (**Figure 6**). When each country actively contributes to the database of SARS-CoV-2 genomes, this protects against sampling biases that can impact the ability to perform phylogenetic analysis and interpret global SARS-CoV-2 data. Towards this goal that affects all of humanity, we advocate for the increased sequencing of SARS-CoV-2 isolates from patients (and infected animals) around the world, and for these data to be shared in as timely a manner as possible.

**Figure 6.**
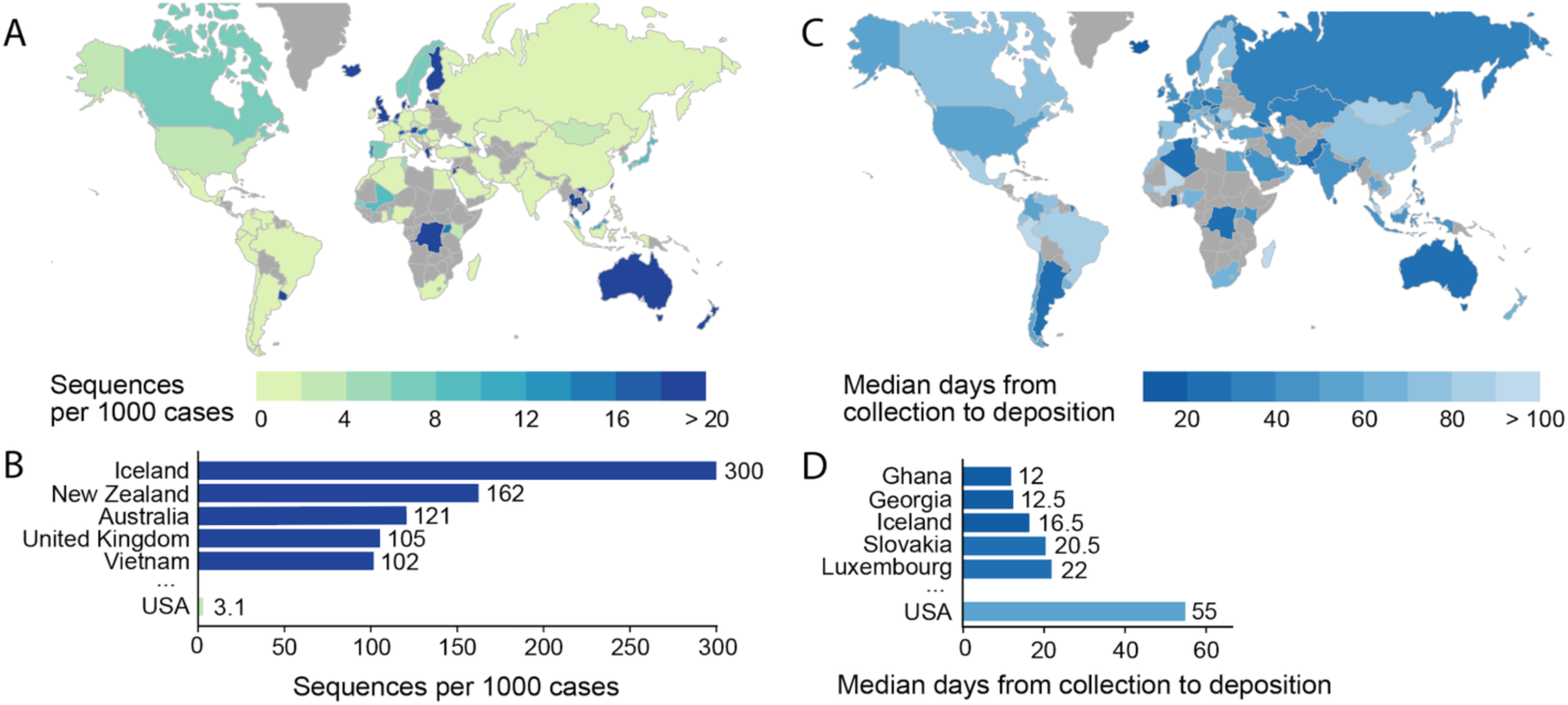
Global statistics of SARS-CoV-2 genomes contribution to GISAID. Interactive and more comprehensive versions of the figure panels are displayed on the Global Sequencing Coverage tab of covidcg.org. **(A)** A world map of countries labeled by the number of SARS-CoV-2 sequences contributed per 1000 cases. **(B)** A bar graph showing the sequences per 1000 cases for the top five countries and the USA. Countries with less than 500 cases were excluded from this plot. **(C)** A world map of countries labeled by median days between sample collection and sequence deposition. **(D)** A bar graph showing the median days from collection to deposition for the top five countries and the USA. These interactive displays are generated using sequencing data from the GISAID EpiCov™ database (nextmeta file) and case data from the JHU CSSE COVID-19 Data (Dong et al., 2020). Only samples that were collected between March and May, 2020 were included to avoid biases from samples that have been collected in the previous three months but not yet deposited onto GISAID.

### Experimental procedures

#### Data Pipeline

Our data processing pipeline is written with the Snakemake scalable bioinformatics workflow engine (Koster and Rahmann, 2012), which modularizes our workflow and enables reproducibility and compatibility with cloud-computing. All code and relevant documentation are hosted on an open-source, publicly available GitHub repository (https://github.com/vector-engineering/COVID19-CG), providing example data for users to validate our pipeline.

#### Data Acquisition

SARS-CoV-2 sequences and metadata are downloaded daily from the GISAID EpiCov™ database (https://epicov.org/epi3/start), by navigating to the “Browse” tab and selecting sequences by “Submission Date”. As of 2020-09-04, only 10,000 sequences can be downloaded from this selection at a time, so the selection is adjusted to include no more than 10,000 sequences. “Sequences”, “Patient status metadata”, and “Sequencing technology metadata” are downloaded separately, stored in separate folders, and renamed for ingestion into the data processing pipeline (see below).

#### Sequence Preprocessing

Based on best practices, we filter out sequences meeting any of the following criteria: (1) Present on the NextStrain’s exclusion list (https://github.com/nextstrain/ncov/blob/master/defaults/exclude.txt), (2) Isolates from non-humans (animals, environmental samples, etc), (3) genome length less than 29,700 nt, or (4) >5% ambiguous base calls. Sequences which pass all preprocessing filters are carried onto the next steps.

#### Metadata Cleaning

We clean metadata with the aim of preserving the original intent of the authors and data submitters while presenting simpler and unified versions to end users. Sequencing metadata is cleaned to remove obvious typos, and to unify labels with the same meaning, e.g., “MinION” and “Nanopore MinION”. Location metadata is cleaned with the goal of simplifying the location selector in the sidebar. Locations with excessive children are collapsed to the nearest upper hierarchical grouping. E.g., if a state has individual data for 200+ towns, these towns will be collapsed to the county level in order to facilitate easier data browsing. Typos and clear identities are also unified to prevent the display of duplicate locations in the application.

#### SNV Assignments

SNVs and insertions/deletions (indels) at the nucleotide and amino acid level are determined by aligning each sequence to the WIV04 reference sequence (WIV04 is a high quality December, 2019 isolate that is 100% identical to the first publicly available SARS-CoV-2 genome reference Wuhan-Hu-1/NC_045512.2, excepting the sequences at the end of the genomes) using *bowtie2*. Spurious SNVs and probable sequencing errors, defined as less than 3 global occurrences, are filtered out prior to downstream analysis. SNVs involving ambiguous base calls (“N” in the original sequences) are ignored. Indels resulting in frameshifts are ignored, and SNVs/indels occurring in non-protein-coding regions are ignored when determining SNVs/indels on the AA level.

#### Lineage/Clade Analysis

Viral lineages, as defined by the *pangolin* tool (Rambaut et al., 2020), and clades (Tang et al., 2020) are provided by GISAID. In accordance with *pangolin*, SNVs present in >90% of sequences within each lineage/clade will be assigned as lineage/clade-defining SNVs.

#### Application Compilation

The web application is written in Javascript, and primarily uses the libraries React.js, MobX, and Vega. The code is compiled into javascript bundles by webpack. All sequence data is compressed and injected inline as JSON into the javascript bundle – no server is needed to serve data to end users. The compiled application files can then be hosted on any static server.

#### Application Deployment

COVID CG (https://covidcg.org) is hosted by Google Cloud Run. The application code is assembled into a Docker image (see Dockerfile), with a build environment (node.js) and deployment environment (NGINX).

## Supporting information

GISAID Acknowledgements

## Acknowledgments

B.D. and A.C. are supported by awards from the National Institute of Neurological Disorders and Stroke (UG3NS111689) and a Brain Initiative award funded through the National Institute of Mental Health (UG3MH120096) and from the Stanley Center for Psychiatric Research. Y.A.C. is supported by a Broad Institute SPARC award and the Stanley Center for Psychiatric Research. We gratefully acknowledge all of the authors from the originating laboratories responsible for obtaining the specimens and the submitting laboratories where genetic sequence data were generated and shared via the GISAID Initiative, on which this resource is based. A full list of authors and contributing laboratories is available (**Supplemental File**).

## Data Availability

All of the data shown in this manuscript and displayed on COVID CG (https://covidcg.org) are downloaded from the GISAID EpiCov™ database (https://www.gisaid.org). All code and relevant documentation are hosted on an open-source, publicly available GitHub repository (https://github.com/vector-engineering/COVID19-CG).

## Author Contributions

Y.A.C., S.H.Z, and A.T.C. conceived the project and browser. B.E.D. supervised the work. A.T.C. and K.A. developed the COVID CG web browser with input from all of the authors. S.H.Z. advised the implementation of lineage and clade analysis. Y.A.C., B.E.D., and A.T.C. prepared the figures, analyzed the data, and wrote the manuscript with input from all authors.

## Supplemental Figures

**Figure S1.**
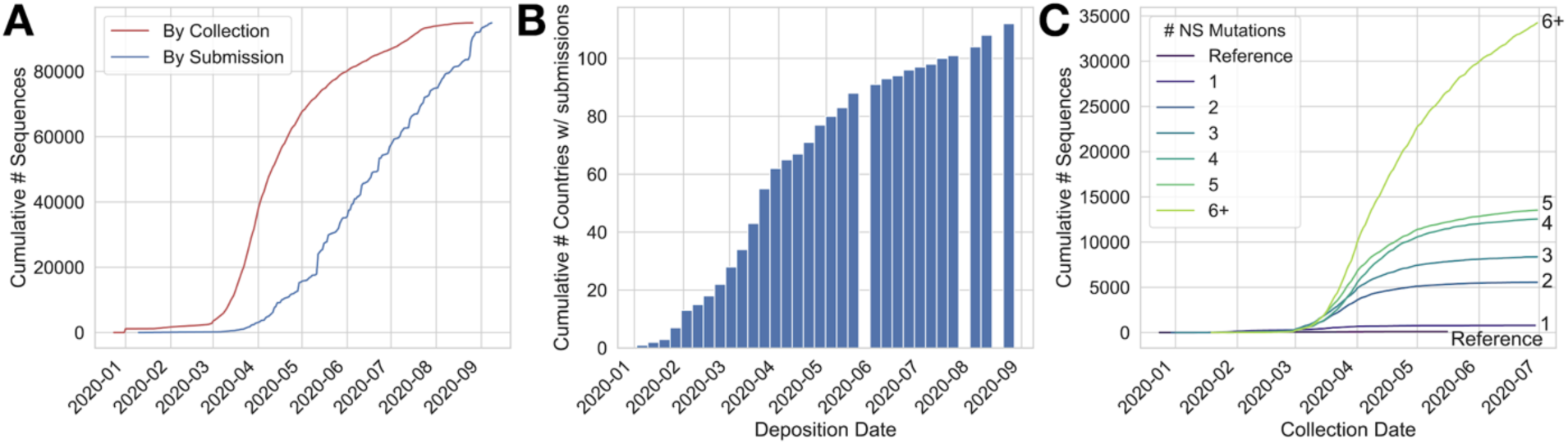
The number of global SARS-CoV-2 genome sequences and mutations is accumulating. Data shown as of September 9, 2020. **(A)** Sequence deposition in GISAID continues at a steady pace, albeit there is a lag between collection (red line) and submission date (blue line). The rate of sequence submission is steady at >10,000 genomes per month. **(B)** More than 100 countries have deposited SARS-CoV-2 genomes in GISAID. **(C)** The number of SARS-CoV-2 variants with more than six nonsynonymous (NS) mutations continues to increase.

**Figure S2.**
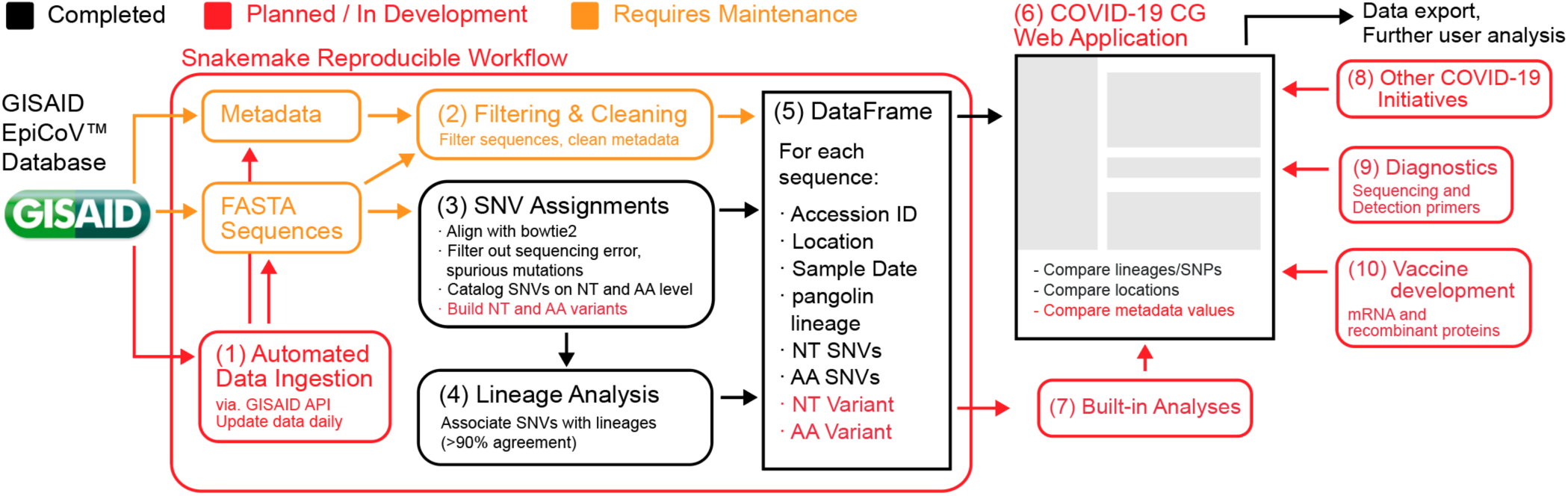
COVID-19 CG computational workflow. **(1)** Starting from the GISAID database, sequences are continuously updated, manually for now, but ultimately via automated data ingestion. **(2)** Based on best practices, we filter out sequences on NextStrain’s exclusion list, non-human isolates, <29,700 nt, or with >5% ambiguous base calls (van Dorp et al., 2020). **(3)** SNVs at the nucleotide and amino acid level are determined by aligning (via *bowtie2*) each sequence to the WIV04 reference, a high quality December, 2019 isolate recommended by GISAID; NextStrain uses the 100% identical Wuhan-Hu-1 (Langmead et al., 2009). Importantly, spurious SNVs and probable sequencing errors are filtered out prior to downstream analysis. **(4)** Viral lineages, defined by the *pangolin* tool, are provided by GISAID. In accordance with *pangolin*, SNVs present in >90% of sequences within each lineage are assigned as lineage-defining SNVs. **(5)** The curated data and metadata, SNVs, and lineage-assigned SNVs are associated with their respective sequence identifier and compiled into a compact data set. **(6)** These data are uploaded onto the COVID-19 CG web application. **(7)** New analyses will be built into the COVID-19 CG application throughout the course of the pandemic. **(8-10)** Features and modules that integrate knowledge from other COVID-19 initiatives are continuously incorporated into COVID-19 CG.

